# Decoding the microbiota of borş: multifunctional potential of a traditional Romanian beverage fermentation

**DOI:** 10.64898/2026.08.13.744363

**Authors:** Silvia-Simona Grosu-Tudor, Annina Meyer, Iulia-Roxana Angelescu, Emanuela-Catalina Ionetic, Ecaterina-Teodora Chirea, Nicholas Bokulich, Stefan Weckx, Luc De Vuyst, Medana Zamfir

**Affiliations:** Institute of Biology of the Romanian Academy, Splaiul Independentei No. 296, Sector 6, 060031 Bucharest, Romania; Laboratory of Food Systems Biotechnology, Department of Health Sciences and Technology, ETH Zurich, Schmelzbergstrasse 7, 8092 Zurich, Switzerland; Research Group of Industrial Microbiology and Food Biotechnology, Faculty of Sciences and Bioengineering Sciences, Pleinlaan 2, B-1050 Brussels, Belgium; FoodFerment, Hoezestraat 65, B-9320 Aalst, Belgium

## Abstract

Romanian borş, a traditional fermented wheat bran beverage, is produced through spontaneous fermentation and represents a complex microbial ecosystem. Despite its cultural importance and presumed health benefits, its microbial ecology and functional potential remain poorly characterized. The present study aimed to elucidate the microbial community structure of borş and link it to functional traits relevant to fermentation performance and food functionality by integrating culture-independent sequencing with culture-dependent isolation and functional characterization. A total of 32 borş samples (12 commercial and 20 homemade) were analyzed. Amplicon-based sequencing revealed a microbiome dominated by lactic acid bacteria (LAB), with lactobacilli accounting for the majority of the bacterial communities and *Lactobacillus amylolyticus* being identified as the most prevalent and abundant species. The yeast communities were mainly composed of fermentative taxa, including *Pichia kudriavzevii* and *Kluyveromyces marxianus*. *Lactobacillus amylolyticus* and *P. kudriavzevii* were also the most frequently isolated species among bacteria and yeasts, respectively. These results highlighted a strong adaptation of the microbial isolates to starch-rich cereal substrates and underscored the central role of these microorganisms in wheat bran fermentation for borş production. Whereas the sequencing-based analyses showed no significant differences in overall diversity between the commercial and homemade borş samples, the cultivation-based results indicated a higher bacterial richness in the commercial products. Notably, the culture-dependent method captured substantially fewer taxa, highlighting the complementary nature of the two approaches. Of a total of 101 bacterial strains (88 LAB and 13 acetic acid bacteria) isolated, many exhibited rapid growth and strong acidification capacity, reaching pH values below 4.5 within 12 h. A functional screening revealed that 21 % of these strains displayed α-amylase activity, 65 % phytase activity, and 50 % β-glucosidase activity, highlighting their capacity to metabolize cereal substrates and enhance the nutrient availability of borş. All strains showed antibacterial activity against at least one indicator bacterium tested, with a universal inhibition of *Listeria monocytogenes*. Overall, Romanian borş harbored a lactic acid bacteria-dominated core microbiome with a significant functional diversity. These findings underscored its potential as a rich source of functional and technologically important strains for application in starter and protective culture development.

## 1. Introduction

Cereal-based fermented beverages have a long tradition, representing one of the oldest forms of food fermentation worldwide [1]. Fermented food products such as borş, boza, kvass, and tarhana are well-recognized examples. They have recently attracted growing scientific and consumer interest because of not only their unique organoleptic characteristics but also their nutritional value and potential health-promoting properties, including micronutrient availability, bioactive compound harboring, enhanced digestibility, antioxidant activity, and probiotic potential [2–4]. Those fermented beverages are still widely produced in Europe and Asia using traditional methods, including spontaneous fermentation and backslopping [5, 6]. Such artisan practices contribute to the development of complex and dynamic microbial ecosystems that shape both product quality and functionality [7].

In general, cereal-based fermented beverages harbor diverse microbial consortia that are represented by lactic acid bacteria (LAB), alongside yeasts and acetic acid bacteria (AAB), which all act together to ensure product preservation and confer nutritional and health-related benefits [2, 4]. Within the group of cereal-based fermented beverages, Romanian borş is a unique souring agent for traditional Romanian soups (*ciorbă*) and a traditional beverage, made through the spontaneous fermentation of wheat bran [8, 9]. Its production is mainly carried out by LAB and yeasts. A preliminary culture-based study investigating the LAB species diversity of borş has reported a prevalence of lactobacilli, among which *Lactobacillus amylolyticus* has been identified as the most prevalent and abundant species [10]. In general, LAB genera, such as *Lactobacillus*, *Leuconostoc*, and *Pediococcus*, are well known to play a central role in similar cereal-based fermentation processes, during which they contribute to acidification, pathogen inhibition, and the development of desirable sensory properties [11].

The recipes for borş manufacture vary considerably, as they are both region-and household-dependent [8, 10]. Nevertheless, the core ingredients remain consistent, encompassing wheat bran that is mixed with warm water and *huşte*, a portion of a previously fermented batch that serves as the natural inoculum. Additional ingredients may include corn flour, beans, aromatic herbs (*e.g.*, lovage or dill), or sour cherry leaves. The fermentation process typically lasts between one and three days, during which the fermented liquor can be harvested daily for immediate consumption, without heat treatment, or it can be replaced with fresh warm water to continue fermentation. Unpasteurized borş that is produced in small-scale artisan facilities is also available in the supermarkets. Its production in these units continues to rely on natural fermentation processes, using parts of previous batches as starter cultures.

The borş fermentation process results in the production of organic acids (mainly lactic acid and acetic acid), trace amounts of ethanol, carbon dioxide, and volatile organic compounds, giving the end-product its characteristic sour taste and slight effervescence [12]. Simultaneously, fermentation enhances the bioavailability of minerals and antioxidants through phytate degradation and an increased release of phenolic compounds [2, 13]. Moreover, the acidic pH (typically 3.5–4.0, sometimes even lower) can suppress the growth of spoilage and pathogenic microorganisms, thus contributing to the microbiological safety and shelf stability of this beverage [14]. Furthermore, borş is believed to be one of the most effective remedies for hangovers, as well as for invigorating the body or fighting anemia (non-scientific literature, such as <u>Fermented Wheat</u> <u>Bran – Romanian Borş</u>, 2014). It is also considered to be beneficial for digestive or respiratory disorders, and for protecting the body against infections (<u>Borşul de casă</u>, 2021; <u>Borşul te scapă</u>, 2015). As these health impacts still have to be proven, borş is a subject of interest for deeper food science and nutritional research.

Despite its widespread traditional use and presumed health benefits, borş remains poorly characterized in the scientific literature, particularly in terms of its microbial ecology and health-related impacts. The spontaneous nature of borş fermentation makes it an interesting subject for microbiological studies, as it provides a natural model for understanding the microbial diversity and/or succession, metabolite production, and in particular strain diversity in non-starter fermentation ecosystems. The latter may lay the basis for the development of tailor-made starter cultures for controlled borş production. Furthermore, a detailed characterization of the microbial and biochemical composition of borş may provide insights into its potential functional properties and contribute to a broader understanding of cereal-based fermented foods and beverages and their bioactive compounds and health-promoting characteristics.

The present study aimed at the investigation of the microbial composition of borş, with particular emphasis on the LAB species involved in the wheat bran fermentation for its production, by using both culture-dependent and culture-independent approaches. Furthermore, the assessment of selected functional traits of isolated LAB strains, including acidification capacity, antimicrobial potential, and enzymatic activities, may contribute to the establishment of possible links between the microbial composition of borş and its functional properties, in turn contributing to the future development of functional starter cultures for healthy borş products.

## 2. Materials and Methods

### 2.1. Collection of borş samples

A total of 32 borş samples, including 12 commercial and 20 homemade products, were collected for this study. They are further referred to as IBB-I-X and IBB-H-X, respectively, with X representing the sample number. Additionally, one sample of *huşte* was collected (IBB-H-008), but results were not included in the present study. Commercial borş batches were produced in Bucharest, Galați, Iaşi, and Buzău, whereas the household samples were collected from local markets located in Botoşani, Suceava, and Bucharest. All samples were collected in sterile vials and immediately stored at-80°C until culture-independent analysis. For the culture-dependent analysis, the samples were stored at 4°C and proccessed within 48-72 h after collection.

### 2.2. Culture-dependent analysis

#### 2.2.1. Microbial enumeration and isolation

For LAB isolation, appropriate serial dilutions of each chilled sample were plated on MRS-5 agar medium [15], supplemented with 0.1 g/l of cycloheximide (Thermo Fisher Scientific, Waltham, Massachusetts, USA) and 0.005 g/l of amphotericin (Apollo Scientific, Manchester, UK) to inhibit fungal growth. The plates were incubated at 30°C for 48 h. Approximately 10 % of the colonies grown on the agar medium were randomly picked, suspended in MRS-5 broth, and purified through succesive passages through both liquid and solid MRS-5 medium. Finally, a total of 288 pure cultures were stored at-80°C in MRS-5 broth, supplemented with 25 % (m/v) glycerol as cryoprotectant.

Yeasts were isolated from nine selected borş samples by plating the serial dilutions on yeast extract-peptone-glucose (YPG) agar medium [10 g/l of peptone (Merck, Darmstadt, Germany), 5 g/l of yeast extract (Merck), 20 g/l of glucose (Merck), 15 g/l of agar (Scharlab, Barcelona, Spain)], supplemented with 0.2 g/l of chloramphenicol (Scharlab) to inhibit bacterial growth. A similar procedure as described above was applied for the isolation, purification, and storage of a total of 81 yeast cultures.

#### 2.2.2. Microbial identification

For microbial identification, DNA was extracted from fresh, overnight cultures by using a Pure Link Genomic DNA kit according to the manufacturer’s protocol (Invitrogen, Carlsbad, California, USA).

To remove redundancies and avoid repetitive analysis, the isolates mentioned above were dereplicated and grouped into clusters applying rep-PCR fingerprinting and numerical cluster analysis. Therefore, (GTG)_5_ primers were used for bacterial dereplicaton, whereas M13 primers were used for yeast dereplication [16]. Representative isolates of each rep-PCR cluster were then identified by sequencing of the 16S rRNA gene (bacteria) and the internal transcribed spacer (ITS) region (fungi) in a commercial facilitiy (Macrogen Europe, Amsterdam, The Netherlands), using the universal primers 27f/1492r [17] and ITS1/ITS4 [18], respectively. The sequence reads obtained were processed and edited using the molecular evolutionary genetics analysis (MEGA) software [19] and identified through comparison with known sequences in the National Center for Biotechnology Information (NCBI) database using the BLAST tool (www.ncbi.nlm.nih.gov/BLAST).

Finally, 101 distinct bacterial strains (88 LAB and 13 AAB) and 22 distinct yeast strains were kept for further analysis. A distinct strain was defined as one representative strain per species per sample.

### 2.3. Culture-independent analysis

#### 2.3.1. DNA extraction, marker gene amplicon library preparation, and sequencing

DNA extraction, amplicon library preparation, and sequencing were performed as described for sourdough previously [20], with minor adaptations for the borş samples. Briefly, 2 mL of borş sample were pelleted (5,000×*g*, 5 min, 4°C), the supernatant was discarded, and the pellet was resuspended in phosphate-buffered-saline (1×PBS) and transferred to a MagMAX™ Bead Beating Plate (Thermo Fisher Scientific). The DNA was extracted using the MagMAX™ Microbiome Ultra Nucleic Acid Isolation Kit on a KingFisher Apex instrument (all from Thermo Fisher Scientific), following the MagMAX Liquid Buccal protocol. ZymoBIOMICS Microbial Community Standard (Zymo Research, Irvine, California, USA) and sterile ultrapure water served as positive and negative extraction controls, respectively. The DNA obtained was stored at −20°C until library preparation.

The bacterial (16S rRNA gene) and fungal (ITS) amplicon libraries were prepared according to the HighALPS ultra-high-throughput barcoding with unique dual indices [21], following the library preparation protocol described previously [20]. The bacterial communities were profiled by targeting the V4 region of the small subunit (SSU) rRNA with the UDI-linked primers 515F and 806R [22, 23]. The fungal communities were profiled via nested PCR of the ITS1 region of the rRNA transcribed unit using the BITS/B58S3 primers [24], followed by barcoding PCR with the UDI-tagged primers. The amplicons obtained were purified with 0.7× Agencourt AMPure XP beads (Beckman Coulter, Brea, California, USA), quantified by a Qubit dsDNA High Sensitivity assay (Thermo Fisher Scientific), pooled at equimolar concentrations, and quality-checked on a high-sensitivity TapeStation.The combined 16S rRNA and ITS libraries were sequenced (300 bp paired-end; 600-cycle kit) on an Illumina NextSeq 2000 with a 20 % PhiX spike-in at the Functional Genomics Center Zürich (Zürich, Switzerland).

#### 2.3.2. Bioinformatic marker gene amplicon processing

The raw paired-end fungal sequences were processed using QIIME 2 (version 2024.10; [25]. The primers and adapters were trimmed with the cutadapt trim-paired plugin [26], followed by denoising via the DADA2 denoise-single plugin [27] with no truncation (--p-trunc-len 0), a maximum expected error of 4.0, and a minimum parent abundance fold-difference of 4.0. The paired-end 16S rRNA-V4 gene sequences were truncated to 150 bp in both directions, and denoised with the DADA2 denoise-paired plugin, with a maximum expected error threshold of 2.0, and a minimum fold-parent-over-abundance threshold of 4.0. For initial non-target filtering, taxonomy was assigned with the classify-sklearn action of the q2-feature-classifier plugin [28] against the customized UNITE v10.99 reference database [29] curated via RESCRIPt [30] for the fungal reads, and the SILVA 138.2 SSU NR99 reference database [31, 32], restricted to EMP 515f/806r amplicon regions prepared using RESCRIPt [30] for the bacterial reads.

For the ITS dataset, the reads assigned to non-fungal sequences, fruiting body–associated taxa, and unclassified phyla, were removed. Subsequently, operational taxonomic units (OTUs) were generated by mapping those filtered fungal reads at 90 % similarity using the VSEARCH cluster-features-closed-reference plugin [33], against the same UNITE v10.99 database, excluding singletons. The OTU features were rarefied to 835 reads per sample, resulting in a final fungal dataset with data for 23 of the 32 collected borş samples. The denoised bacterial reads were filtered to exclude non-bacterial sequences (*e.g.*, mitochondria, chloroplasts, archaea, and eukaryotes) and subsequently clustered into OTUs at 97 % identity using the VSEARCH cluster-features-closed-reference plugin against the SILVA 138.2 database. The resulting OTU feature table was rarefied to 465 reads per sample, resulting in a final dataset of 26 of the 32 borş samples.

#### 2.3.3. Statistics

The alpha-diversity (richness, evenness, and Shannon entropy) and beta-diversity (Jaccard distance and Bray–Curtis dissimilarity) metrics were calculated using the q2-diversity plugin in QIIME 2 [25]. The beta diversity estimates were calculated based on the rarefied OTU features. The alpha-diversity metrics of the household and commercial samples were compared using two-sided Mann-Whitney U tests, and the *p*-values were adjusted with the Bonferroni-Holm false discovery rate (FDR) multiple-testing correction across the different metrics [34]. The beta-diversity was visualized by using a principal coordinate analysis (PCoA), with confidence ellipses added at 95 %. Global PERMANOVAs comparing household samples to commercial samples were performed in scikit-bio [35–37]. To align the culture-dependent to the culture-independent microbial community data, a Procrustes analysis was performed with n = 1000 permutations to evaluate the statistical significance of the correspondence of the beta-diversity metrics between the two approaches [38]. A simple linear regression modeling was used to compare the alpha-diversity metrics between the culture-dependent and-independent data, split by household and commercial borş samples, without inclusion of fixed or random effects [39].

To compute differential abundances between household and commercial borş samples, ANCOM-BC2 wrapped into the composition ancombc2 plugin was applied [40], with structural zeros enabled and a prevalence cut-off of 0.05, and the *p*-values were adjusted with Benjamini Hochberg-FDR [34].

Sankey visuals were generated in plotly [41] for joint-visualization of the abundant and prevalent taxonomic clades. The weighting (of the branch widths) was based on a simple, composite prevalence-abundance index. For each taxon *i*, the prevalence was computed as the fraction of samples with relative abundance > 10^−4^, and mean relative abundance across all samples, *r̅_i_*. Then, the log of the averaged relative abundance ℓ*_i_* was defined according to Equation 1, both the mean prevalence (Equation 2) and ℓ*_i_* (Equation 3) were scaled by min–max scaling across taxa, and a core score was defined as the product of the scaled prevalence and scaled log-mean abundance (Equation 4).

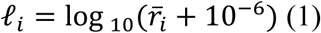

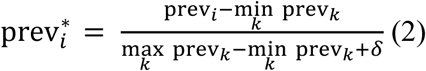

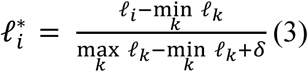

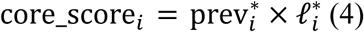

Sparse partial-correlation networks were computed as previously described (Meyer et al., 2026), by using Graphical Lasso with cross-validation [42, 43]. Non-zero entries in the resulting precision matrix defined network edges, feature identity (bacteria, fungi) was used for node coloring and the above defined core score for node size (Equation 4). Topological metrics - degree centrality, betweenness centrality, closeness centrality, and eigenvector centrality - were calculated to mathematically identify key taxa in each of the networks.

All the statistical and computational analyses of the microbiome data were performed in Python (v3.10.14), unless stated otherwise. The data preprocessing used pandas [44] and NumPy [45]. The dimensionality reduction and network cross-validation was performed with scikit-learn [43]. The statistical modeling (Mann-Whitney) used statsmodels [46]. The visualizations were generated with Seaborn [47] and Matplotlib [48], plotly was used for Sankey visuals [41], and NetworkX (v2.8.8) for microbial networks [49].

### 2.4. Functional properties analysis of distinct bacterial strains

#### 2.4.1. Growth and acidification in MRS-5 medium

For the growth analysis, each LAB strain was inoculated (2 %, v/v) in 200 µL of MRS-5 broth in sterile 96-well microtiter plates, in triplicate. The microplates were incubated at 30°C, and the optical density at 600 nm (OD_600_) readings were recorded every hour for 24 h, using a FLUOstar Omega microplate reader (BMG-Labtech, Ortenberg, Germany) equipped with temperature control and shaking function.

For the acidification analysis, the fastest growing strains were inoculated in 20 mL of MRS-5 broth and incubated at 30°C for 24 h. Samples were collected at 0, 3, 6, 12, and 24 h to measure the pH value with an InoLab 720 pH meter (WTW, Weilheim, Germany).

#### 2.4.2. Enzymatic activities

The α-amylase, phytase, and β-glucosidase activities of the LAB and AAB strains grown in MRS-5 medium were assessed using plate-based or spectrophotometric assays.

The α-amylase production was screened for on starch agar medium composed of 1.0 % (m/v) soluble starch (Merck), 0.3 % beef extract (Merck), and 1.5 % agar (Scharlab), adjusted to pH 7.5. Overnight bacterial cultures were spotted (10 µL) on the surface of the agar medium and the plates were incubated at 30°C for 96 h. Following this incubation, the agar surfaces were flooded with iodine solution, composed of 1.0 % iodine (Merck) and 2.0 % potassium iodide (Merck). The formation of clear halos around the spots indicated starch hydrolysis.

The phytase activity of the LAB and AAB strains was measured using a colorimetric assay based on the release of inorganic phosphate [50]. Briefly, overnight cultures were centrifuged (8,000 x *g*, 10 min, 4°C), the cell pellets were washed with 50 mM phosphate buffer (pH 7.0) and then resuspended in sterile ultrapure water to obtain an OD_600_ value of about 0.6 at a 1:10 dilution. These suspensions were incubated with 3 mM sodium phytate (BLD Pharmatech, Shanghai, China) in 0.2 M sodium acetate (Merck) of pH 4.0 at 45°C for 1 h. The reaction was stopped with 5.0 % trichloroacetic acid (Merck), and liberated phosphate was determined by staining with ammonium molybdate (Lach-Ner, Neratovice, Czech Republic) and ferrous sulphate (Merck), and absorbance reading at 700 nm. One unit of phytase activity corresponded to an increase in absorbance of 0.001 U per min.

The β-glucosidase activity was determined using p-nitrophenol-β-D-glucopyranoside (pNPG; Thermo Fisher Scientific) as the substrate [51]. Therefore, washed LAB cell pellets were incubated with 2.5 mM pNPG at 40°C for 30 min. The reaction was terminated by incubation at 95°C, and the release of p-nitrophenol was quantified by measuring the absorbance at 410 nm against ultrapure water. One unit of β-glucosidase activity was defined as the amount of enzyme liberating 1 μmol of p-nitrophenol per mL per min under the assay conditions.

#### 2.4.3. Antibacterial activity

The antibacterial activity of the LAB and AAB strains was evaluated using the agar well diffusion method [52], with slight modifications related to the sample preparation protocol. Overnight cultures of the strains were centrifuged (8000 x *g*, 10 min, 4°C), and the resulting cell-free supernatants were concentrated tenfold by freeze-drying. The indicator strains used in this assay were *Staphylococcus aureus* ATCC 25923, *Escherichia coli* ATCC 25922, *Listeria monocytogenes* ATCC 1911-1, *Salmonella enterica* ATCC 14028, *Bacillus cereus* CBAB, *Bacillus subtilis* ATCC 6633, and *Lactobacillus delbrueckii* subsp*. bulgaricus* LMG 6901^T^. The latter strain was inoculated in a soft MRS agar overlay (0.7 % agar, m/v), which was poured over a basic MRS agar layer (Oxoid, Basingstoke, Hampshire, UK), whereas the other indicator strains were spread directly onto a BHI agar medium (Oxoid) using sterile cotton swabs. Afterward, wells of approximately 8 mm in diameter were cut in the agar media and the holes were filled with the concentrated supernatants. The plates were kept at room temperature for 30 min to allow the supernatants to diffuse into the agar medium prior to incubation at the optimum growth temperature of the indicator strains (28°C for the two *Bacillus* strains, and 37°C for all the other strains tested). The inhibition halos around the wells were measured after 24 h of incubation.

#### 2.4.4. Statistics

When applicable, the results of the growth parameters and functional properties of the LAB and AAB strains tested were statistically analyzed with GraphPad Prism (GraphPad Software, Boston, Massachusetts, USA).

## 3. Results

### 3.1. Isolation and taxonomic identification of bacteria and yeasts present in household and commercial borş samples

Except for three household borş samples, bacterial colonies could be recovered from the MRS-5 agar media for 17 household samples and 12 commercial samples. Among the 101 distinct strains of the 288 colonies isolated, purified, and dereplicated, the 88 strains identified as LAB through the 16S rRNA gene sequencing protocol belonged to 26 different species (Fig. 1A). The five most frequently isolated LAB species across the borş samples analyzed were *L. amylolyticus*, *Lacticaseibacillus paracasei*, *Limosilactobacillus fermentum*, *Lentilactobacillus buchneri*, and *Furfurilactobacillus rossiae* (Fig. 1A). Among those species, *L. amylolyticus* was isolated from ten of the 17 household samples and from four of the 12 commercial samples. This was followed by *L. paracasei*, isolated from ten samples (seven commercial), and *L. fermentum*, recovered from nine samples (five homemade). *Lentilactobacillus buchneri* and *F. rossiae* were isolated predominantly from commercial borş samples. Most of the remaining taxa (21 species) were recovered only occasionally, occurring in one or two products. Those species included *Lacticaseibacillus casei, Lactobacillus helveticus, Leuconostoc citreum*, and *Schleiferilactobacillus perolens* (Fig. 1A). Also, 13 strains were identified as AAB, belonging to eight different *Acetobacter* species. *Acetobacter lovaniensis* was isolated from both household and commercial borş samples at similar frequencies (three samples each). It was the fifth most common bacterial taxon isolated from all samples.

**Fig. 1.**
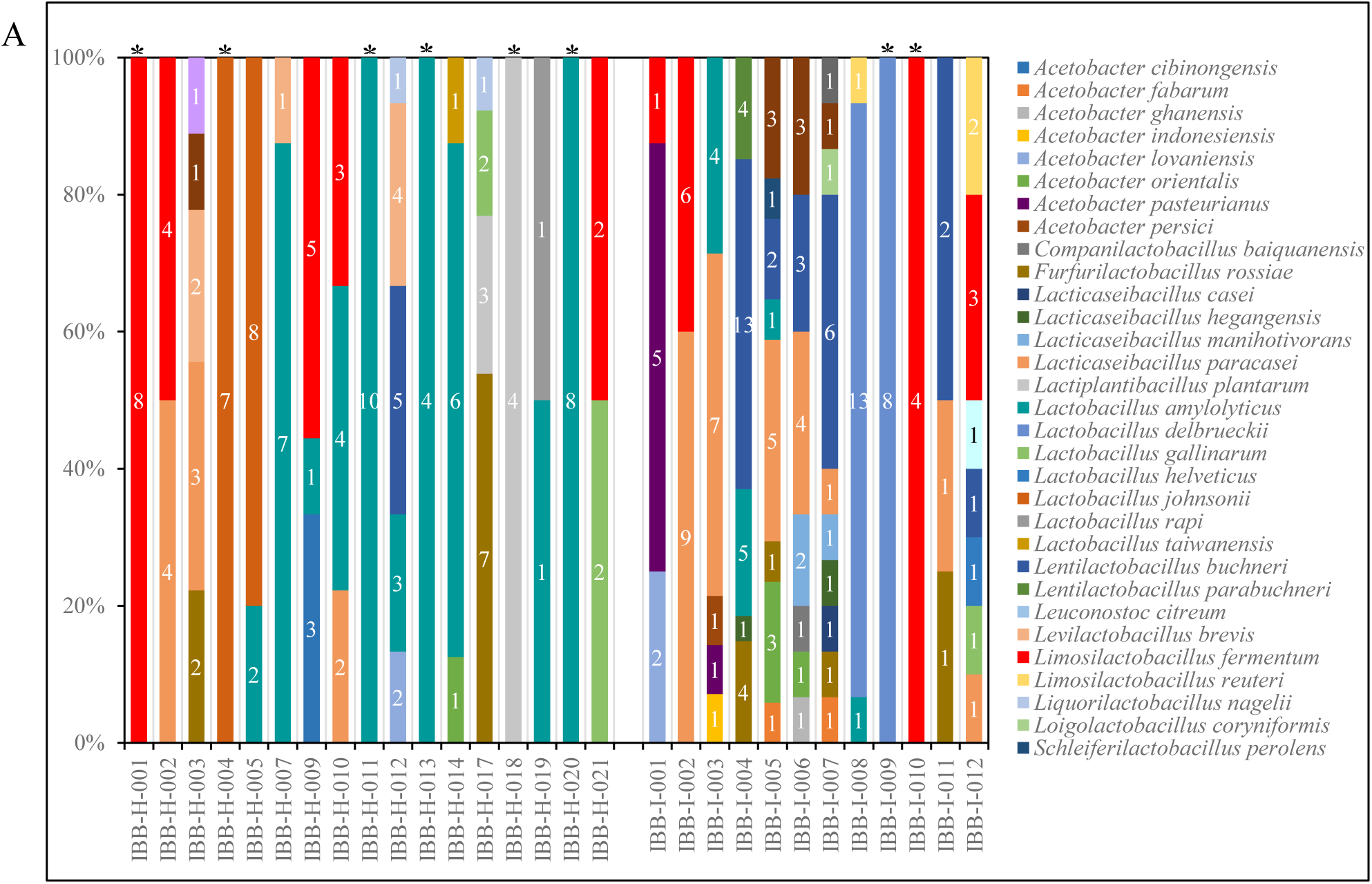
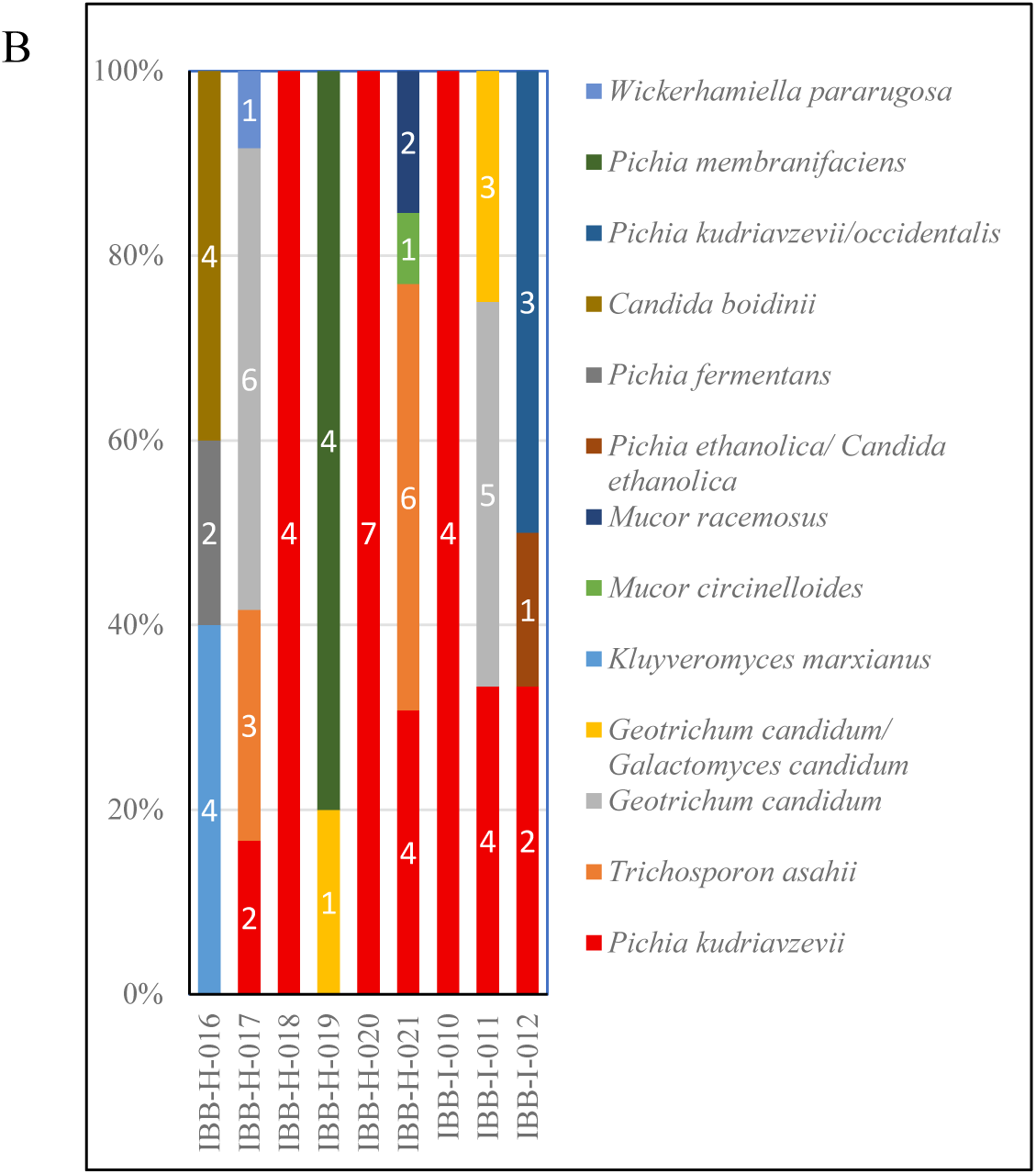
Microbial diversity of Romanian borş samples determined by a culture-dependent approach. (A) Distribution of bacterial species identified across the borş samples analyzed. The number of isolates for each taxon is shown on the corresponding bar. Samples containing only a single isolated species are marked with an asterisk. The first group of samples are homemade (IBB-H), and the second group are commercial samples (IBB-I). (B) Distribution of yeast species identified across the borş samples analyzed. The number of isolates for each taxon is shown on the corresponding bar.

The distribution of all these bacterial species over the 17 household and 12 commercial borş samples showed that the LAB and AAB species diversity varied among the samples, ranging from one to ten different bacterial species per sample. Overall, the commercial borş samples tended to recover more individual bacterial species per single sample, with the highest number of different species isolated from samples IBB-I-007 (10 species), IBB-I-005 (8 species), and IBB-I-006 and IBB-I-012 (7 species each). In contrast, the lowest bacterial diversity, represented by only one single species, occurred in six household and only two commercial borş samples (indicated by an asterisk in Fig. 1A). From an additional five borş samples, four of which were homemade, only two bacterial species were isolated and identified.

Among the yeasts isolated and identified, the prevailing species was *Pichia kudriavzevii*, recovered from seven of the nine selected borş samples. Other *Pichia* species, such as *Pichia fermentans* and *Pichia membranifaciens*, as well as *Geotrichum candidum* and *Trichosporon asahii* were isolated from one or two samples (Fig. 1B).

### 3.2. Culture-independent identification of bacteria and yeasts present in household and commercial borş samples

Culture-independent, amplicon-based sequencing revealed the presence of LAB in all 26 bacterial datasets and yeasts in all 23 fungal datasets (post rarefaction). AAB were additionally identified in 11 of the 26 bacterial datasets. Across both household and commercially produced borş samples, the predominant LAB, AAB, and yeast species were *L. amylolyticus*, *A. lovaniensis*, and *P. kudriavzevii*, respectively.

### 3.3. Culture-independent and culture-dependent community diversity across all borş samples

Concerning bacteria, the culture-independent β-diversity did not differ significantly between the household and commercial borş samples (PERMANOVA on Bray–Curtis and Jaccard distances; FDR-P > 0.05; Fig. 2A,B). In contrast, the culture-dependent profiles showed an origin-specific structuring; both the Bray–Curtis (R² = 0.094, FDR-P = 0.009) and Jaccard (R² = 0.078, FDR-P = 0.010) distances differed significantly between the household and commercial samples (Fig. 2C,D). For both approaches, the commercial borş samples exhibited significantly higher bacterial richness (Fig. 2E,F; FDR-P < 0.05), whereas the Shannon diversity and evenness did not differ between origins.

**Fig. 2.**
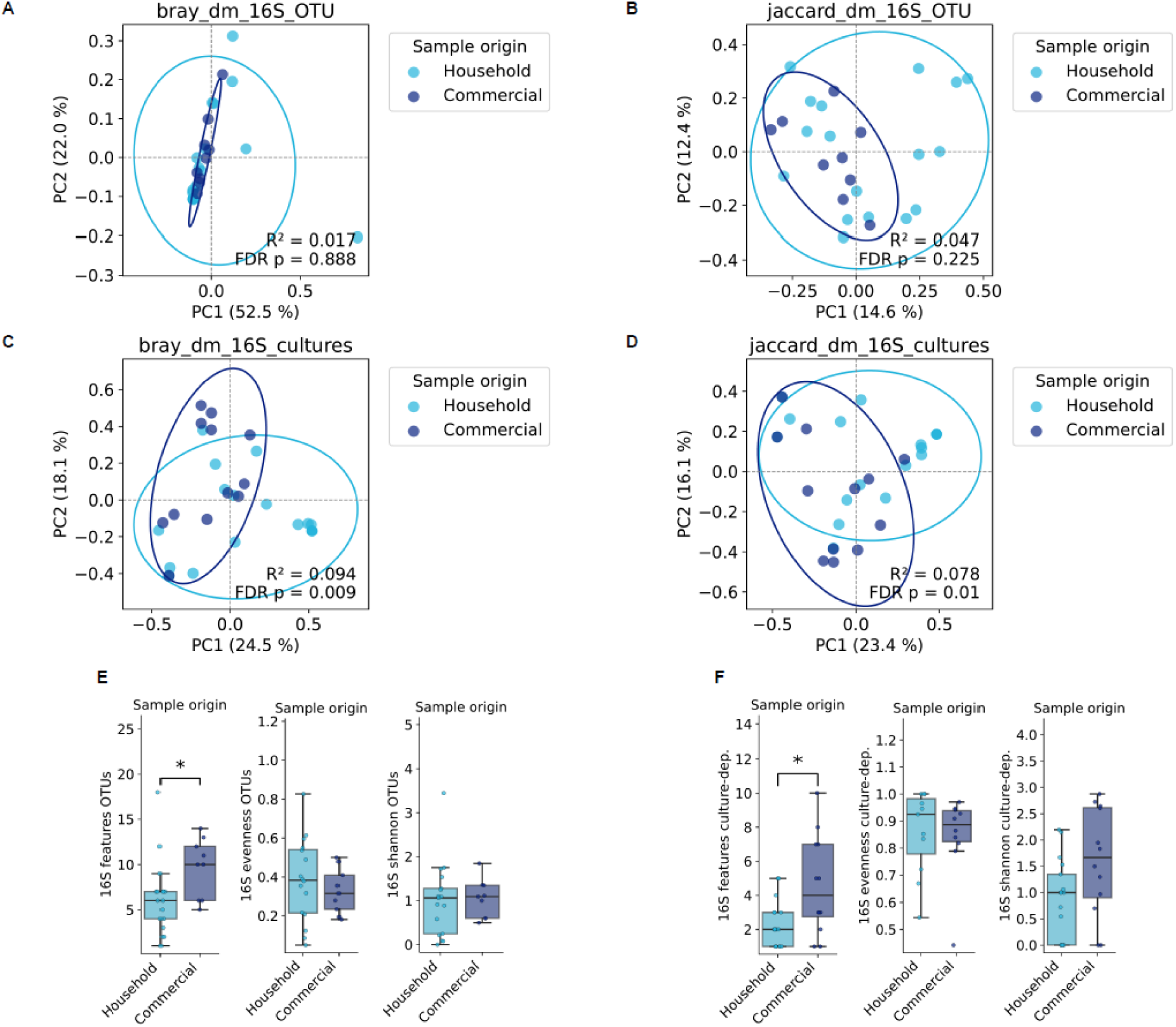
Comparison of the culture-independent and culture-dependent bacterial community diversity across Romanian household and commerical borş samples. (A,B) PCoA plots of Bray–Curtis (A) and Jaccard (B) distances based on culture-independent (16S v4 rRNA-based sequencing) bacterial community profiles. (C,D) Corresponding PCoA plots for culture-dependent bacterial diversity. Ellipses indicate 95 % confidence intervals; the PERMANOVA results comparing the household and commercial bors samples are shown for each distance metric (BH–FDR–corrected p values). (E,F) Bacterial alpha-diversity (richness, evenness, Shannon index) of household versus commercial bors samples for (E) sequencing-based and (F) culture-dependent community data. Pairwise differences were assessed using Mann–Whitney U tests; p values were Benjamini–Hochberg FDR–corrected, and asterisks indicate adjusted p < 0.05. For fungi, neither the β-diversity nor the α-diversity differed significantly between the household and commercial borş samples, for either the sequencing-based or the culture-based data (Fig. 3A–F).

**Fig. 3.**
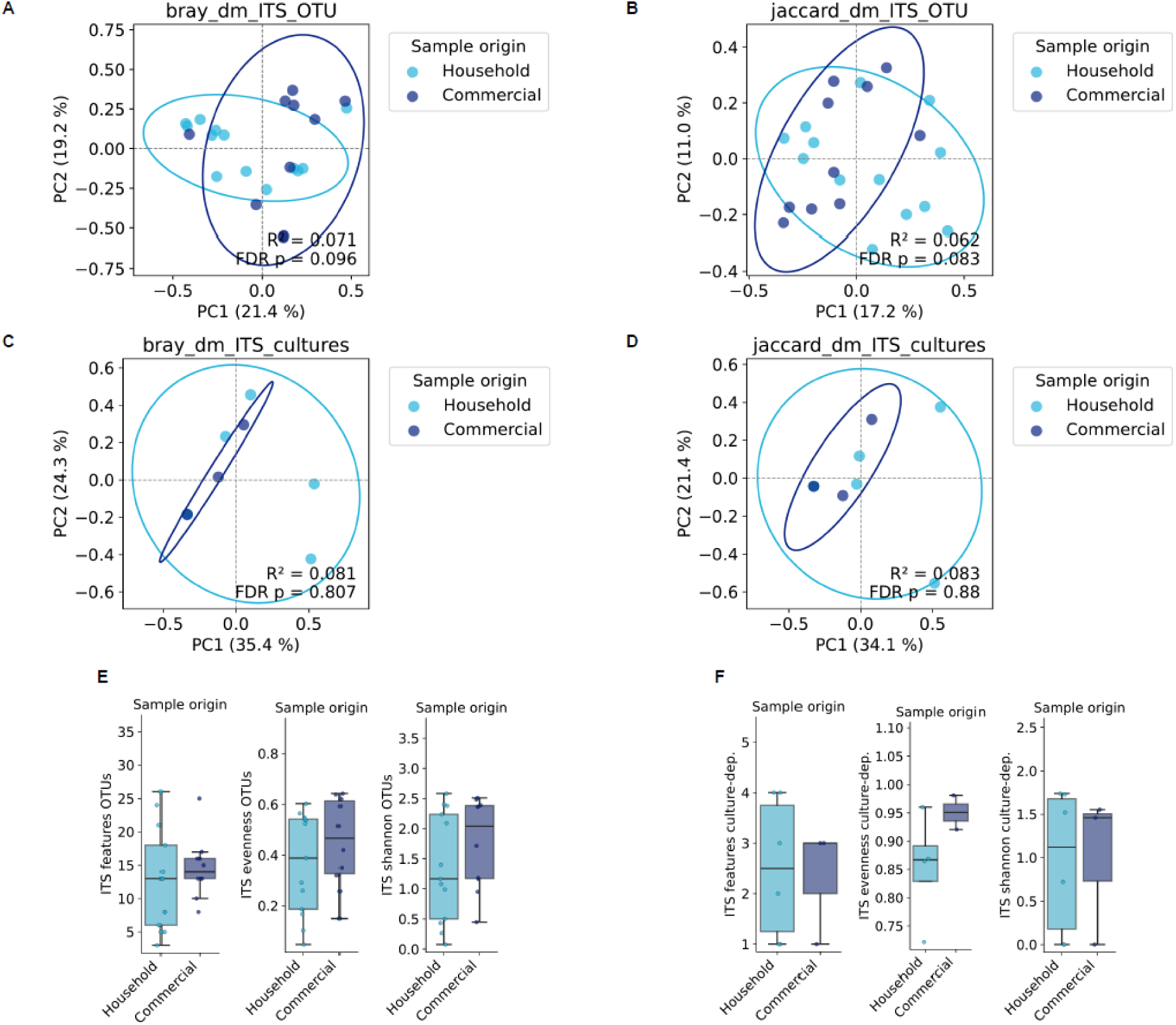
Comparison of the culture-independent and culture-dependent fungal community diversity across Romanian household and commerical borş samples. (A,B) PCoA plots of Bray–Curtis (A) and Jaccard (B) distances based on culture-independent (ITS1-based sequencing) fungal community profiles. (C,D) Corresponding PCoA plots for culture-dependent fungal diversity. Ellipses indicate 95 % confidence intervals; the PERMANOVA results comparing household and commercial borş samples are shown for each distance metric (BH–FDR–corrected *p* values). (E,F) Fungal alpha-diversity (richness, evenness, Shannon index) in household versus commercial bors samples for (E) sequencing-based and (F) culture-dependent community data. Pairwise differences were assessed using Mann–Whitney U tests; *p* values were Benjamini–Hochberg FDR–corrected, and asterisks indicate adjusted *p* < 0.05.

The concordance between the culture-independent and culture-dependent data was limited, with a low correspondence of β-diversity feature spaces for bacteria (Jaccard, M² = 0.67; Bray–Curtis, M² = 0.92) and low to moderate correspondence for fungi (Jaccard, M² = 0.37; Bray–Curtis, M² = 0.32). On average, the culture-dependent approach recovered approximately five-fold fewer bacterial taxa per sample than the culture-independent amplicon-based sequencing method, estimated by a simple linear regression modelling.

### 3.4. Core microbial clades and differentially abundant taxa in the borş microbiomes

To identify clades that are both prevalent and abundant - and thus likely most relevant to borş production - a core score was applied to the borş microbiome data (Equation 4), combining the scaled mean relative abundance and prevalence across all samples. This score was subsequently used to weight the bacterial and fungal taxa in Sankey visualisations (Fig. 4A,B).

**Fig. 4.**
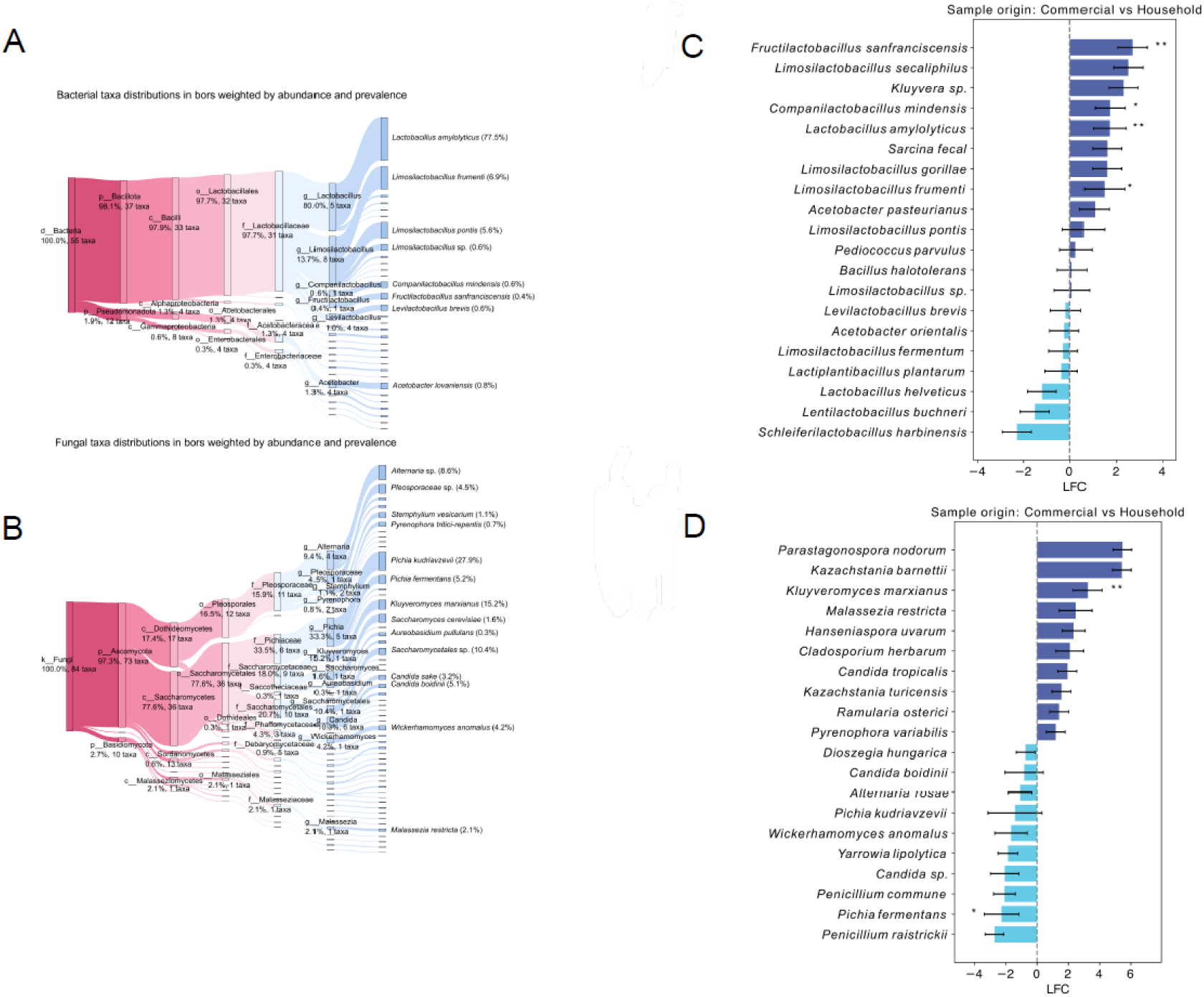
Importance of microbial clades and taxa in wheat bran fermentation for Romanian borş production. (A,B) Sankey plots visualising the most frequent and abundant bacterial (A) and fungal (B) clades in the bors samples analyzed. Branch width is proportional to the core score combining both prevalence and abundance of all taxa in the branch (Equation 4); thus, thicker branches indicate clades that are both more prevalent and more abundant. Key nodes are labelled with the mean total relative abundance of the branch across all borş samples (percentage) and the number of distinct taxa detected within that branch. (C,D) Differentially abundant bacterial (C) and fungal (D) taxa between household and commercial borş samples, inferred by ANCOM-BC2. Bars show estimated log-fold changes (LFCs) with standard deviations; taxa with Benjamini–Hochberg FDR–corrected *p* values < 0.05 (*) or < 0.01 (**) are indicated.

Among the bacterial communities, species of the LAB genus *Lactobacillus* prevailed in the core microbiome (Fig. 4A). The lactobacilli accounted for 80.0 % of the total relative abundance (five taxa), with *L. amylolyticus* being the single most impactful and abundant species (total relative abundance of 77.5 %). Species of the genus *Limosilactobacillus* contributed 13.7 % of the total relative abundance (eight taxa), with *Limosilactobacillus frumenti* (total relative abundance 6.9 %) as the most abundant representative. The fungal core clades were predominated by the class of the Saccharomycetes (total relative abundance of 77.6 %, 36 taxa) and the class of the Dothideomycetes (17.4 %, 17 taxa) (Fig. 4B). The former clade comprised fermentative yeasts, including *P. kudriavzevii* (total relative abundance 27.9 %), *Kluyveromyces marxianus* (15.2 %), *Candida sake* (3.2 %), *Candida boidinii* (5.1 %), and *Wickerhamomyces anomalus* (4.2 %), whereas Dothideomycetes were mainly represented by plant-associated genera, such as *Alternaria*, *Pleosporaceae*, *Stemphylium*, and *Pyrenophora*.

A differential-abundance analysis (ANCOM-BC2) revealed distinct origin-specific bacterial signatures (Fig. 4C). *Fructilactobacillus sanfranciscensis*, *Companilactobacillus mindensis*, *L. amylolyticus*, and *L. frumenti* were all significantly more abundant in the commercial than in the household borş fermentations, with estimated differences of approximately 1–2 log-fold changes (FDR-*P* < 0.05). For the fungi, *K. marxianus* was significantly enriched in the commercial borş fermentations, whereas *P. fermentans* was more abundant in the household ones, with effect sizes of 3 and 2 log-fold changes, respectively (Fig. 4D).

### 3.5. Microbial network architecture and stability of core taxa

To examine how the core taxa are embedded within their communities in the different production environments, separate graphical Lasso networks for the household and commercial borş samples were inferred, each including the 20 highest core-score taxa (bacteria and fungi combined) and their neighbours (Fig. 5).

**Fig 5.**
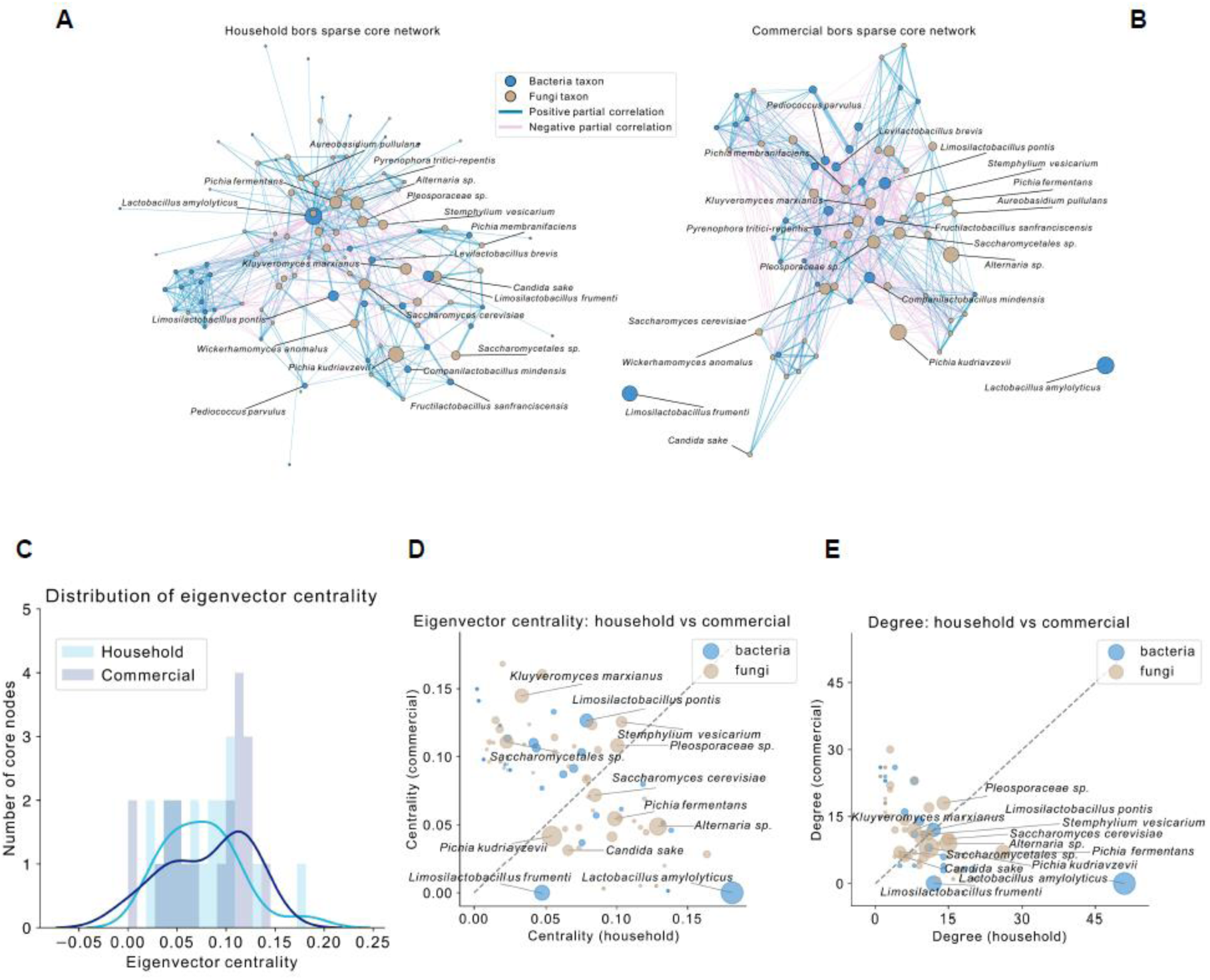
Variation of the Romanian borş core-microbiome across production origins. (A,B) Sparse partial-correlation networks inferred using graphical Lasso for household (A) and commercial (B) borş samples. Each network includes the 20 taxa with the highest core scores (bacteria and fungi combined) and their directly connected neighbours. The node size is proportional to the core score (prevalence–abundance score), the node colour indicates bacterial versus fungal origin, and the edges represent positive or negative partial correlations; the edge width reflects the absolute partial correlation strength. Core taxa (top 20) are labelled. (C) Distributions of eigenvector centrality for the household and commercial borş networks. (D) Eigenvector centrality in the household borş network plotted against that in the commercial borş network for each taxon; the point size indicates the core score, and the core taxa are labelled. (E) Degree centrality of taxa in household and commercial borş networks, with bubble size again proportional to the core score and core taxa labelled.

Both networks were organized into several modules, with most core taxa occupying the main central module. Notably, the two most abundant and impactful LAB species, *L. amylolyticus* and *L. frumenti*, were central within the household borş network (Fig. 5A), but in the commercial borş network they remained prevalent and abundant while becoming largely disconnected from the rest of the network (Fig. 5B). Overall, the node centrality tended to be higher in the commercial borş network (Fig. 5C), consistent with the denser connectivity. A comparison of the eigenvector centrality between the networks showed marked shifts for the key LAB species, particularly *L. amylolyticus* and *L. frumenti*, whereas core yeast species, such as *P. kudriavzevii, P. fermentans, Saccharomyces cerevisiae*, and *Candida* spp., displayed a more stable centrality across both origins (Fig. 5D). The degree centrality similarly differed between the household and commercial borş networks, again highlighting the origin-specific changes in connectivity for the core LAB species (Fig. 5E).

### 3.6. Growth and acidification curves of distinct LAB strains in MRS-5 medium

Most of the bacterial strains selected displayed good growth in MRS-5 medium, reaching OD_600_ values between 0.85 and 1.66 after 24 h of incubation (Fig. 6A). A few strains demonstrated notably faster growth compared to the others. For example, *Lactiplantibacillus plantarum* HF082 reached a high OD_600_ value within the first 6 h of incubation, which continued to increase upon further incubation. Other fast-growing strains included *L. fermentum* HF299 and two *Levilactobacillus brevis* strains, namely, HF085 and HF231.

**Fig. 6.**
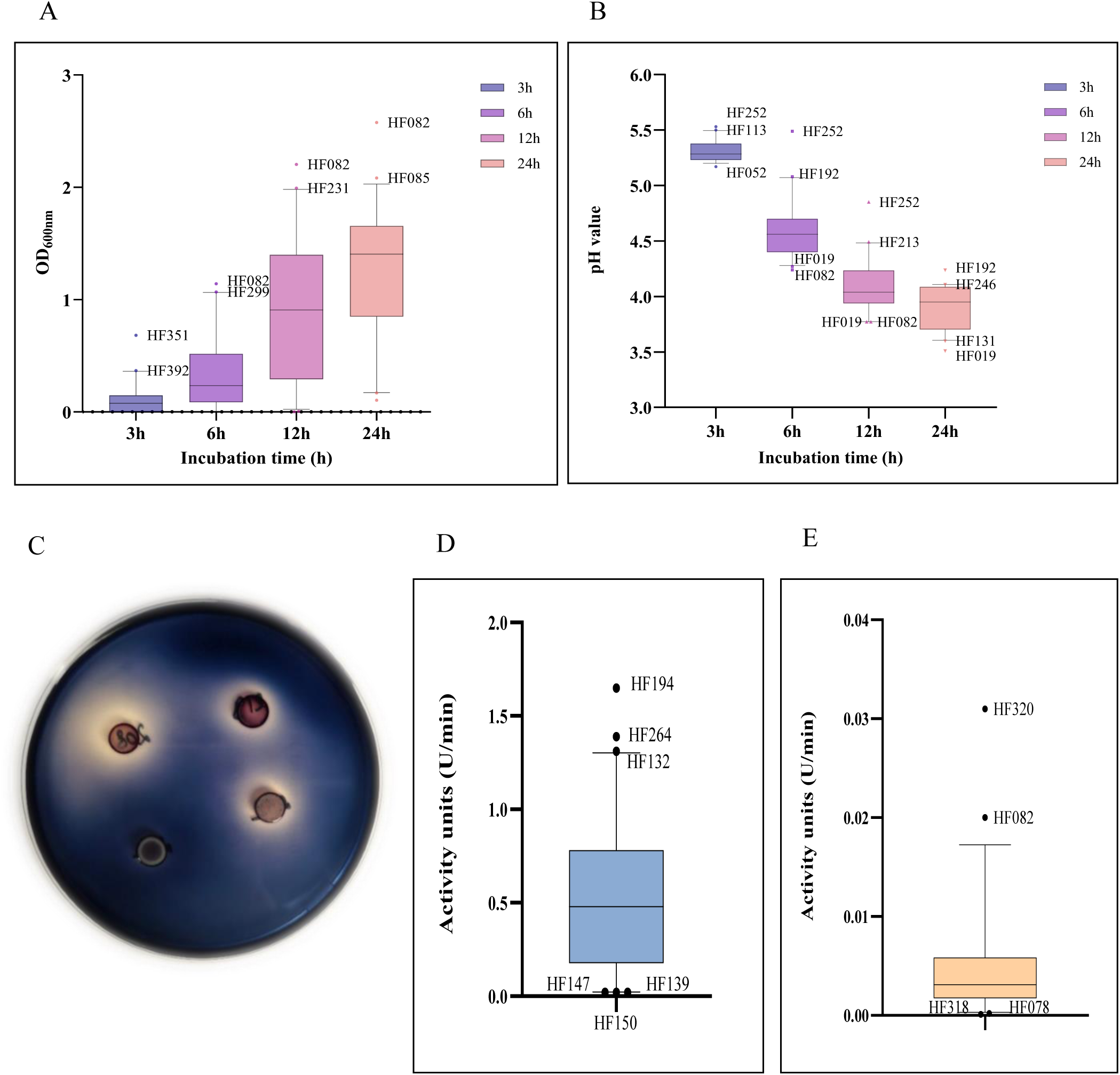
Functional characterization of selected bacterial strains from Romanian borş. (A) Growth of lactic acid bacteria (LAB) strains in MRS-5 medium at 30°C, presented as box plots (2.5–97.5 %) of optical density (OD_600_) measurements. Outlier strains exhibiting the fastest growth are highlighted in the graph. (B) pH reduction over time of the bacterial cultures tested, shown as box plots (10–90 %). Outlier strains exhibiting faster/slower acidification are indicated in the graph. (C) Amylase production by LAB strains isolated from borş samples. Positive strains are indicated by the presence of a clear halo surrounding the bacterial colony after staining with iodine solution. (D) Box plots (5–95 %) showing the phytase activity of bacterial strains grown in MRS-5 medium. Outlier strains are indicated on the graph and labeled with their corresponding strain codes. (E) Box plots (5–95 %) showing the β-glucosidase activity of bacterial strains grown in MRS-5 medium. Outlier strains are indicated on the graph and labeled with their corresponding strain codes.

Twenty of the fastest growing strains acidified the MRS-5 medium to a pH of 4.4-4.7 within the first 6 h of incubation (Fig. 6B). The median pH value reached 4.6 after 6 h of incubation and decreased to 4.0 after 12 h. *Lactiplantibacillus plantarum* HF082 and *Lactiplantibacillus argentoratensis* HF019 displayed the most rapid acidification (pH 4.2 after 6 h of incubation). Strain HF019 produced the most acidic culture overall, reaching a pH of 3.5 after 24 h of incubation.

### 3.7. Enzymatic activities of the distinct LAB strains

Of 101 bacterial strains tested (both LAB and AAB), 21 showed a positive reaction for α-amylase activity (Fig. 6C; Table 1). Most of these strains were identified as *L. fermentum*, *L. amylolyticus*, and *L paracasei*.

**Table 1.** Distribution of α-amylase positive strains among the bacterial species isolated from Romanian borş samples.

| Bacterial species | α-amylase positive strains | Total number of strains tested |
| --- | --- | --- |
| *Lactobacillus amylolyticus* | 5 | 14 |
| *Lacticaseibacillus paracasei* | 4 | 11 |
| *Limosilactobacillus fermentum* | 7 | 9 |
| *Lentilactobacillus buchneri* | 3 | 8 |
| *Acetobacter indonesiensis* | 1 | 1 |
| *Acetobacter orientalis* | 1 | 3 |
| Other LAB species | 0 | 46 |
| Other AAB species | 0 | 9 |
| Total | 21 | 101 |

Sixty-six strains displayed phytase activity when grown in MRS-5 medium. The median enzymatic activity was 0.48 U/min, with most values ranging between 0.18 and 0.78 U/min (Fig. 6D). The highest phytase activities were recorded for *L. delbrueckii* HF194 (1.65 U/min), *L. plantarum* HF264 (1.39 U/min), and *L. paracasei* HF132 (1.31 U/min).

Only 50 strains exhibited β-glucosidase activity, in general between 0.002 and 0.006 U/min, with a median of 0.003 U/min (Fig. 6E). *Lactobacillus nagellii* HF320 (0.03 U/min) and *L. plantarum* HF082 (0.02 U/min) showed the highest activities.

### 3.8. Antibacterial activity

All 101 strains tested were able to inhibit the growth of at least one of the indicator strains used in the agar well diffusion assay. Notably, the majority of the strains exhibited antibacterial activity against multiple indicators (Fig. 7A), with 28 strains (26 LAB and 2 AAB strains) showing inhibition against all indicator strains tested. An additional 47 strains inhibited between two and four indicator strains, whereas only 13 strains displayed a narrow inhibitory spectrum, affecting just a single indicator strain.

**Fig. 7.**
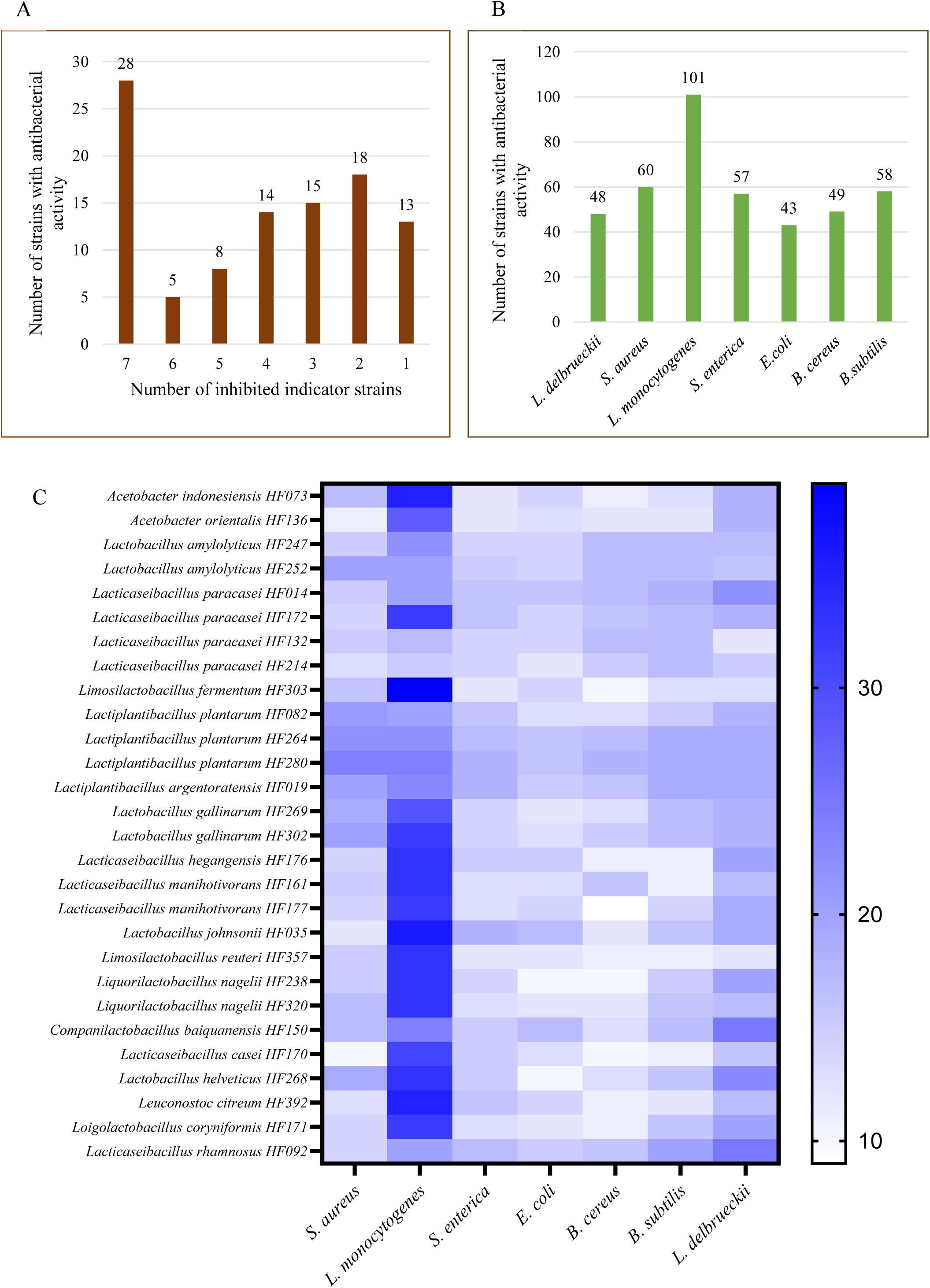
Antibacterial activity of lactic acid bacteria and acetic acid bacteria strains isolated from Romanian borş samples. (A) Inhibitory spectrum of the bacterial strains. Numbers above each bar represent the number of strains that were able to inhibit the corresponding number of indicator strains. (B) Susceptibility of the indicator strains to the antibacterial effect of borş-derived bacterial strains. The number of strains that were able to inhibit each indicator strain is indicated on each bar. (C) Heatmap showing the diameters of the inhibition zone that indicate the antibacterial activity of selected borş-derived strains.

*Listeria monocytogenes* ATCC 1911-1 was by far the most sensitive indicator strain, its growth being inhibited by all strains tested (Fig. 7B). In contrast, *E. coli* ATCC 25922, *L. delbrueckii* subsp. *bulgaricus* LMG 6901^T^, and *B. cereus* CBAB were less susceptible to inhibition, since only 43, 48, and 49 strains, respectively, had an antibacterial activity against them. *Staphylococcus aureus* ATCC 25923, *S. enterica* ATCC 14028, and *B. subtilis* ATCC 6633 had a similar susceptibility, all being inhibited by 58-60 strains (Fig. 7B).

Among the 28 strains with broad inhibitory spectrum (active against all indicator strains), four strains belonged to the LAB species *L. paracasei*, and three strains to the LAB species *L. plantarum* (Fig. 7C). Other LAB and AAB species were also found in this group, but less well represented (by one or two strains).

## 4. Discussion

The present study, which was built on previous work that first reported the bacterial composition of Romanian borş [10], provided a comprehensive characterization of the Romanian borş microbiota by analyzing 20 household and 12 commercial borş samples. Integrating taxonomic profiling with functional analyses enabled decoding the roles of these microorganisms in the context of fermentation performance, nutritional enhancement, and sensory development. Since a culture-dependent diversity analysis is inherently constrained by cell viability during transport/storage and isolation conditions, this borş study combined culture-dependent and culture-independent approaches to characterize the borş microbiome and directly compare the diversities captured by sequencing of whole communities *versus* recovered isolates.

With respect to the culture-dependent approach, the absence of bacterial colonies on MRS-5 agar medium in three household borş samples may be attributed to the absence of live cells or microbial death occurring either during borş storage or transport of the borş samples to and their storage in the laboratory [20]. Homemade borş is typically unpasteurized and stored in closed bottles or large containers, usually in cool environments, but without strict temperature or aeration control, all of which may influence the viability and laboratory recovery of the microbial cells.

The use of MRS-5 agar medium for bacterial isolation yielded a greater microbial diversity for both the household and commercial borş samples compared to the use of a standard MRS medium [53] in the previous study [10]. This enhanced diversity may be attributed to the enriched composition of MRS-5 medium, which includes multiple carbon sources and a vitamin mixture, supporting the growth of LAB strains with high nutritional demands [54]. Nevertheless, the bacterial diversity recorded per sample by cultivation was still about five times lower than that obtained by culture-independent DNA sequencing.

As expected, LAB were the most represented bacteria for borş, as shown by both culture-dependent (88 of 101 strains identified and belonging to 26 different species) and culture-independent methods. This predominance indicated a successful wheat bran fermentation process, with an essential role of these LAB strains in acidification (via production of lactic acid), flavor development (carbohydrate and amino acid metabolisms), and preservation (low pH) [55]. The growth of AAB (13 strains belonging to 8 species of the *Acetobacter* genus) was likely favored by the low initial pH of the medium and the aerobic incubation conditions [56]. AAB are recognized for their ability to produce acetic acid, in turn causing acidification, and various flavor-active compounds, thereby playing a significant role in shaping the sensory profile of the final product [57].

Among the five prevailing LAB species identified culture-dependently, *L. amylolyticus*, *L. fermentum*, and *L. paracasei* were the most frequent. As *L. amylolyticus* also had the highest prevalence and abundance in the culture-independent analysis, it may be a key functional LAB species during wheat bran fermentation for borş production, being well adapted to the starch-rich environment characteristic for this fermentation process. The species *L. amylolyticus* was first described in beer malt and wort, in which it displays amylolytic activity [58]. It was further found in other fermented food products, such as wheat sourdough or tofu whey [59]. Its amylolytic activity has been linked to the presence of starch-degrading enzymes, such as amylase and pullulanase [60, 61]. The fact that *L. amylolyticus* could not be isolated from all borş samples in which it was detected culture-independently, may indicate that MRS-5 medium, lacking starch as (additional) energy source, is not the optimal medium for its isolation. However, the prevalence of *L. amylolyticus* was significantly higher in the commercial borş samples analyzed culture-independently, whereas isolation on MRS-5 agar medium showed a higher incidence of this taxon in the household samples. This again indicates a loss of cell viability during storage or transport, or the absence of appropriate substrates (*e.g.*, starch) or specific growth factors in the isolation medium. Alternatively, the occurrence of *L. amylolyticus* colonies may have been overlooked in samples with high bacterial diversity, which was the case for certain commercial borş samples. Other prevailing LAB species, such as *L. fermentum* and *L. paracasei*, were frequently isolated from MRS-5 agar medium, although very low abundant (*L. fermentum*) or not detected (*L. paracasei*) by V4-16S rRNA gene sequencing. In contrast, *L. frumenti* and *L. pontis* were only detected by 16S rRNA gene sequencing and were more abundant and prevalent in the household borş samples. These species are known to request specific nutrients and growth factors, a reason why MRS-5 medium was introduced for their isolation from sourdough [54]. *Limosilactobacillus fermentum* is associated with fermented vegetables [62], naturally fermented cheeses [63], and traditional cereal-based fermentation processes, such as sourdough, trahanas, or boza [64]. In these products, *L. fermentum* may enhance their preservation, improve their sensory attributes, and contribute to their nutritional value, among other beneficial properties [65]. *Lacticaseibacillus paracasei* is commonly associated with dairy products, but it is also frequently detected in plant-based fermented beverages, kefir grains, and kombucha [66]. This LAB species has been extensively studied owing to its significant commercial relevance and potential health-promoting properties [67, 68]. *Limosilactobacillus frumenti* and *L. pontis* are well-adapted to cereal matrices and are commonly recovered from sourdough and other traditional cereal-based fermentation processes, including fermented cereal beverages and backslopped products, in which diverse LAB communities persist [69, 70]. Their metabolic activities contribute to rapid acidification, the development of distinct sensory profiles (through organic acid and volatile organic compound production), and various nutritional improvements (*e.g.*, improved digestibility and bioavailability of nutrients) in fermented cereal foods [2, 71].

As nine LAB and six AAB species were only isolated from one borş sample, and always along with other LAB or AAB species, they may represent transient or opportunistic microorganisms introduced via the raw materials or as a result of environmental exposure. However, these species may be responsible for the spoilage of borş, as is the case for *S. perolens*, a known soft drink spoilage bacterium [72]. Nevertheless, some of these rare species may also contribute beneficially to a fermented food product, for example, *Leuc. citreum*, which can produce mannitol, antimicrobial compounds, and dextran [73].

Although the commercial borş samples generally displayed a higher microbial diversity than the homemade ones, none of the commercial producers indicated the use of a defined starter culture, meaning that their wheat bran fermentation processes likely rely on backslopping, similar to the approach used for homemade borş. This higher diversity in the commercial borş samples could be linked to the controlled conditions of industrial fermentation processes, such as a constant temperature, pH monitoring, and hygiene practices, all of which can support the stable coexistence of multiple microbial taxa [74]. In contrast, homemade fermentation processes are inherently more variable and influenced by environmental fluctuations, which may favor the prevalence of a few robust strains. Additionally, the microbial composition of homemade borş may also reflect the specific ingredients used, the artisan manufacturing practices applied, and the handling procedures employed, which often vary not only by region but also from household to household, as is the case for homemade sourdoughs [75–77]. The origin-specific structuring of the culture-dependent bacterial diversity of the household and commercial borş samples elucidated that three commercial borş samples (IBB-I-005, IBB-I-006, and IBB-I-007) exhibited the highest bacterial diversity, with seven to ten different taxa identified in each. These three products originated from the same producer, located in Bucharest, and shared between four and six bacterial species, highlighting the role of controlled environmental conditions in maintaining a stable core microbiota. The differences in microbial composition among those samples may be attributed to variations in ingredients, such as the use of lovage in IBB-I-005 and celery in IBB-I-006.

Regarding the yeast communities of borş, the culture-independently identified core taxa included not only *P. kudriavzevii*, *K. marxianus*, and *Candida* species, but also some plant-associated genera, albeit in a much lower relative abundance. As in other related cereal-based fermented food products, yeasts may contribute to the fermentation process by metabolizing simple saccharides released from the cereal starch, and they may enhance the aromatic complexity of the final products by generating a wide range of volatile organic compounds [78]. They may also contribute to improved nutritional properties by facilitating the degradation of antinutritional factors and enhancing nutrient bioavailability [2]. *Pichia kudriavzevii* was also the predominant species isolated on YPG agar medium, proving its importance in wheat bran fermentation for borş production. The safety of this yeast species has been a subject of debate for a long time, especially due to its resistance to fluconazole [79]. However, *P. kudriavzevii* frequently occurs in traditional fermented foods and beverages worldwide, including fermented cereals, and this yeast species gained a lot of interest in recent years because of its potential applications in food processing and biotechnology [80]. Whereas it has occasionally been associated with spoilage, *P. kudriavzevii* more often contributes positively to flavor development and nutritional quality, and some strains even show probiotic potential [81, 82]. The yeast diversity did not correlate with the origin of the samples, albeit that the *Pichia* and *Wickerhamomyces* species were more abundant in the household borş samples, whereas the *Kazachstania* and *Kluyveromyces* species were predominantly associated with the commercial borş samples. In addition, both the bacterial and yeast communities included taxa that were unique to a single production origin. This might be the result of differences in the raw materials used, processing conditions applied, and microbial sources encountered between the household and commercial borş productions. Certain bacterial and yeast taxa may become specific to one production origin, reflecting their adaptation to those particular fermentation environments [7].

Altogether, the culture-dependent and culture-independent data revealed a LAB-dominated core microbiome for Romanian borş, with origin-specific peripheral bacterial and fungal taxa. Interestingly, representative isolates from these core and accessory clades performed differently in terms of growth, acidification, enzymatic activities, and antibacterial potential. This underlined their specific roles in wheat bran fermentation for borş production. Most of the 101 bacterial isolates tested exhibited good growth in MRS-5 medium, indicating that this medium supports the proliferation of a broad range of strains. Additionally, the strong acidification potential of 20 selected LAB strains, with the majority reducing the pH to below 4.5 within the first 12 h of growth, suggested a high level of metabolic activity and efficient carbohydrate fermentation, consistent with desirable functional traits for the development of starter cultures [83]. From a technological perspective, *L. plantarum* HF082 and *L. argentoratensis* HF019 appeared particularly promising, exhibiting the most rapid acidification among all strains tested (reaching a pH of approximately 4.2 within only 6 h of incubation). A rapid reduction of the pH is a critical trait of starter cultures, as it enhances the fermentation safety of the raw materials by suppressing the growth of spoilage and pathogenic microorganisms, and it contributes to desirable flavor development in the final product [84]. Moreover, *L. argentoratensis* HF019 achieved the lowest final pH, as low as 3.5, indicating not only a highly active metabolism but also strong acid tolerance, which is an advantageous feature for maintaining dominance throughout a fermentation process [85].

A further characterization of distinct bacterial strains included the assessment of their enzymatic activities, specifically those of α-amylase, phytase, and β-glucosidase, which play key roles in the breakdown of complex carbohydrates, mineral release, or enhancement of nutritional value during cereal fermentation [86]. A total of twenty-one strains, among which *L. fermentum*, *L. amylolyticus*, and *L. paracasei,* demonstrated α-amylase activity, as shown by starch-iodine staining, indicating their ability to hydrolyze starch and suggesting an enzymatic adaptation to carbohydrate-rich environments. As these species were also among the most frequently isolated ones from the borş samples examined, the idea that amylase production is a key functional trait in cereal-based fermentation processes, such as wheat bran fermentation for borş production, was supported, because starch serves as a primary energy source in this matrix. Hence, these bacteria likely initiated starch degradation, releasing simple saccharides that can support their own growth or that of other fermentative microorganisms [87]. Amylolytic LAB capable of utilizing starch as their sole energy source are relatively rare and are found within just four genera [86]. *Lactobacillus amylovorus* was one of the first LAB species described for its amylolytic activity [88], but subsequent studies have also confirmed starch-degrading capabilities for *L. amylolyticus*, *L. fermentum*, and *L. paracasei*, among others [86]. The fact that approximately 65 % of the strains tested demonstrated phytase activity, whereas around 50 % exhibited β-glucosidase activity, indicated a widespread enzymatic potential for phytate degradation, and for the hydrolysis of flavonoids and phenolic compounds, respectively. Strains exhibiting strong phytase activity, such as *L. delbrueckii* HF194, *L. plantarum* HF264, and *L. paracasei* HF132, may enhance the mineral bioavailability and simultaneously reduce antinutritional factors in borş, thereby improving the overall nutritional quality of this fermented beverage [89]. Strains such as *L. nagelii* HF320 and *L. plantarum* HF082 that displayed high β-glucosidase activity likely contributed to the breakdown of complex molecules and the development of flavor compounds, as well as to the release of health-promoting bioactive compounds during wheat bran fermentation [90].

Finally, all LAB strains tested demonstrated inhibitory activity against *L. monocytogenes* ATCC 1911, which is consistent with the well-documented ability of LAB to combat Gram-positive pathogens through the production of organic acids and bacteriocins [91]. In contrast, *E. coli* ATCC 25922 was the least susceptible, reflecting the characteristic resilience of Gram-negative bacteria to bacteriocins and acidic conditions owing to their protective outer membrane [92]. These findings were in line with previous reports indicating that antimicrobials produced by LAB tend to be more effective against Gram-positive bacteria than Gram-negative ones [93]. Moreover, the inhibitory activity of AAB strains against the pathogens used in the present study was likely caused by the production of acetic acid [94, 95]. The widespread antimicrobial activity among the strains tested suggested the production of broad-spectrum antimicrobial compounds, such as organic acids, or synergistic actions of inhibitory metabolites [96]. Strains capable of inhibiting both Gram-positive and Gram-negative pathogens may serve as promising bioprotective cultures for the production of fermented foods and beverages, contributing to an enhanced microbial safety and prolonged shelf-life [97]. The narrow inhibitory spectrum of eleven LAB strains, limited to only one or two indicator strains, suggested the production of more target-specific antimicrobials, such as certain bacteriocins [98]. Overall, the antibacterial activity data showed the great potential of borş-derived LAB and AAB strains as members for the development of starter cultures with protective functions, or probiotic cultures [99].

## 5. Conclusions

The present study provided the most comprehensive characterization to date of the Romanian borş microbiome, expanding earlier findings by integrating culture-independent sequencing with extensive culture-dependent isolation and functional profiling. By directly comparing these complementary approaches, this study demonstrated that while amplicon-based sequencing captures a substantially broader microbial diversity, cultivation remained indispensable for linking taxonomic identity to technological and functional traits relevant to fermentation, safety, and nutritional quality of a fermented food product.

Among the LAB-dominated core, the recurrent detection and high abundance of *L. amylolyticus* highlighted its strong adaptation to starch-rich cereal substrates and underscored its central role in wheat bran fermentation for borş production.

A functional screening revealed a high technological potential among borş-derived isolates. Many bacterial strains exhibited rapid growth and strong acidification capacity, efficient starch degradation, phytate hydrolysis, β-glucosidase activity, and broad-spectrum antibacterial effects. Taken together, the physiological and functional analyses of the bacterial isolates demonstrated that the microbial communities associated with Romanian borş represent a valuable reservoir of strains with multifunctional capabilities, combining efficient fermentation performance, enzymatic versatility, and antimicrobial activity. These advantages make them promising candidates for the development of defined, starter or protective cultures aimed at improving the quality, safety, and consistency of borş and other traditional fermented beverages.

## Acknowledgements

The authors acknowledge financial support from the project HealthFerm, which is co-funded by the European Union under the Horizon Europe grant agreement No. 101060247 and the Swiss State Secretariat for Education, Research and Innovation (SERI) under contract No. 22.00210. Views and opinions expressed are however those of the author(s) only and do not necessarily reflect those of the European Union or European Research Executive Agency (REA). Neither the European Union nor REA can be held responsible for them. Part of the work was financially supported by the research project no. RO1567-IBB11/2026 from the Institute of Biology Bucharest of the Romanian Academy. E.-T. Chirea and E.-C. Ionetic are PhD students with a scholarship at the School of Advanced Studies of the Romanian Academy-SCOSAAR. The authors thank Michelle Neugebauer for technical support for microbiome sequencing and the Genomic Diversity Center of ETH Zurich for their support with amplicon library preparation.

## Author contributions

MZ, LDV, NB, conceived and designed the experiments. SSGT, AM, IRA, ECI, and ETC performed the experiments. AM - bioinformatic analysis of microbiome sequencing data including conceptualization, methodology, formal investigation and visualization. SSGT, IRA, ECI, and ETC – culture-dependent analysis, screening for functional properties. MZ, AM, and LDV wrote the manuscript. NB and SW contributed to the revision of the manuscript. All authors read and approved the paper. MZ, NB, SW – funding acquisition.

## Data availability

All sequence data will be deposited on EBI-ENA upon acceptance of the manuscript. Source data (metadata) along with taxa abundance tables have been deposited together with code notebooks in GitHub (see code availability).

## Code availability

Code notebooks for bioinformatic processing, statistical analyses, network generation, and visualizations have been deposited and are openly accessible on GitHub at https://github.com/anninameyer/bors-microbiome-analysis.

